# An FPGA-based hardware accelerator supporting sensitive sequence homology filtering with profile hidden Markov models

**DOI:** 10.1101/2023.09.20.558701

**Authors:** Tim Anderson, Travis J. Wheeler

## Abstract

**Background:** Sequence alignment lies at the heart of genome sequence annotation. While the BLAST suite of alignment tools has long held an important role in alignment-based sequence database search, greater sensitivity is achieved through the use of profile hidden Markov models (pHMMs). The Forward algorithm that provides much of pHMMs’ sensitivity is relatively slow, motivating extensive efforts to increase speed. Numerous researchers have devised methods to improve pHMM alignment speed using hardware accelerators such as graphics processing units (GPUs) and field programmable gate arrays (FPGAs). Here, we describe an FPGA hardware accelerator for a key bottleneck step in the analysis pipeline employed by the popular pHMM aligment tool, HMMER.

HMMER accelerates pHMM Forward alignment by screening most sequence with a series of filters that rapidly approximate the result of computing full Forward alignment. The first of these filters, the Single Segment ungapped Viterbi (SSV) algorithm, is designed to filter out 98% of non-related inputs and accounts for ~70% of the overall runtime of the DNA search tool *nhmmer* in common use cases. SSV is an ideal target for hardware acceleration due to its limited data dependency structure.

**Results:** We present Hardware Accelerated single segment Viterbi Additional Coprocessor (HAVAC), an FPGA-based hardware accelerator for the SSV algorithm. The core HAVAC kernel calculates the SSV matrix at 1739 GCUPS on a Xilinx Alveo U50 FPGA accelerator card, ∼ 227x faster than the optimized SSV implementation in *nhmmer*. Accounting for PCI-e data transfer data processing, HAVAC is 65x faster than nhmmer’s SSV with one thread and 35x faster than nhmmer with four threads, and uses ∼ 31% the energy of a traditional high end Intel CPU. Because these computations are performed on a co-processor, the host CPU remain free to simultaneously compute downstream pHMM alignment and later post-processing.

**Author summary:** Sequence alignment lies at the heart of genome sequence annotation, and must be both fast and accurate. Signals of relationships between sequences are obscured over time by mutational forces, so that alignment and annotation of the full diversity of life demands highly sensitive tools. Profile hidden Markov models (pHMMs) provide the greatest sensitivity in the face of diversity, but are relatively slow. Here, we describe an approach to improving the speed of pHMM search that leverages field programmable gate arrays - hardware devices that can be configured to implement arbitrary digital circuits to achieve impressive parallelism and energy efficiency. Our tool, HAVAC, accelerates one key bottleneck step in the analysis pipeline employed by the popular pHMM aligment tool, HMMER. HAVAC produces a ∼ 60x speedup over the analogous stage in HMMER. HAVAC can be implemented as a part of a larger sequence homology search tool for faster search times and reduced energy usage. Interested users can download HAVAC on github at https://github.com/TravisWheelerLab/HAVAC.

## Introduction

After assembling the genome of an organism, it is standard practice to annotate the contents of that genome by comparing it to a library of known sequences. When the organism is evolutionarily distant from other sequenced genomes, as is common in the context of environmental metagenomic samples, high quality annotation depends on maximizing sensitivity in that comparative analysis. To date, high sensitivity in sequence comparison is achieved through sequence alignment, in which the letters of two sequences are arranged to identify regions of similarity. In the context of sequence alignment, models of mutational probability are used to compute a measure of the significance of the resulting alignment. Here, we focus on alignment methods for sequences that are highly divergent; see Sahlin et al. [1] for a review of methods for rapidly matching nearly identical sequences (as in the context of read mapping [2–4]).

Sequence alignment has been the target of intense design advances for algorithmic and statistical inference methods over several decades, resulting in sensitive and remarkably fast approaches for sequence annotation. For many years, the dominant tool in the space of high-volume sequence alignment was BLAST [5], with support from carefully-designed scoring models like position specific scoring matrices [6, 7]. In the decades since the introduction of BLAST, advances in sensitivity have come primarily in the form of position specific scoring matrices [8–10] and eventually profile hidden Markov models (pHMMs) [11]. Thanks to robust strategies for training and scoring [12, 13] and their representation of position-specific probabilities of observing letters, insertions, and deletions, pHMMs have remained the state-of-art for sensitive sequence annotation [14, 15].

Orthogonal to this development of sensitive models has been an ever-present push for greater speed, motivated by the exponential growth of modern sequence databases. These speed gains are generally achieved by either (i) filtering candidate alignments data with less computationally expensive algorithms, or (ii) devising faster implementations of high sensitivity alignment algorithms. The most popular alignment-based annotation tools achieve their speed by the first strategy, avoiding data analysis through various fast methods for predicting whether a sequence has the potential to produce a high score when exposed to a relatively expensive alignment algorithm [5, 16–20]. These approaches typically depend on indexing either the target sequences, query sequences, or both, and using the resulting indices to identify promising “seeds” for more intensive processing.

Any work-avoidance strategy runs the risk of lost sensitivity due to avoiding candidates containing true positive matches. In sequence alignment, index-based seed finding methods address these sensitivity/speed trade-offs through careful selection of data structures and parameterization. While recent advances retain BLAST-like sensitivity with 30-100x speed gains [21], the sensitivity of full-featured pHMMs is still unrivaled.

The alternative acceleration strategy (*apply the same core algorithm, but faster*) typically depends on some form of hardware acceleration. One such strategy leverages the SIMD (Single instruction, multiple data) vector instructions available on all modern CPUs [22]. SIMD sequence alignment implementations [23–26] have achieved impressive speed gains. This technique serves as a core part of the acceleration strategy used in popular pHMM alignment tools [21, 27, 28].

Hardware acceleration for sequence alignment has also been developed in the context of specialized hardware such as graphics processing units (GPUs [29]) and field programmable gate arrays (FPGAs [30]). Here, we introduce an FPGA-base hardware accelerator that speeds up the key bottleneck stage of the HMMER pipeline by as much as 60x. Before presenting detailed methods and results, we first provide a brief introduction to the relevant aspects of HMMER, followed by a light introduction to hardware acceleration.

## The HMMER Pipeline

The high sensitivity of pHMMs is due to (i) position-specific probabilities learned from multiple members of a sequence family, and (ii) application of the Forward algorithm [31] for computing support for the relationship between query HMM and target sequence. Computation of a sequence alignment amounts to discovering a path through a *Q*x*T* 2-dimensional matrix, where Q and T are the lengths of the query model and target sequence respectively. In the context of pHMMs, the Viterbi algorithm identifies a most-probable path through through that matrix, and computes support for the relationship between Q and T from the single corresponding alignment. This is functionally equivalent [32] to the Smith-Waterman algorithm [33] that are approximated by BLAST and other faster tools mentioned above. Meanwhile, the Forward algorithm computes the sum of the probabilities of all possible paths (all alignments), and uses this as the basis for measuring support for relatedness. Both algorithms show run time complexity Θ(*QT*) [34], but Forward is much slower than Viterbi due to increased constant run time factors [35].

HMMER3 [35] produced a *>*100x speedup over the prior release, thanks to development of a pipeline consisting of faster pHMM alignment filters that approximate the Forward score. The key idea is that the Forward algorithm is only applied to a small number of candidate sequence matches that are allowed to pass these earlier, faster filters. Specifically: query/target pairs are aligned using a SIMD vectorized [23, 25, 36] implementation of the Viterbi algorithm, using reduced precision 16-bit integers in place of floating point scores; this Viterbi stage approximates the score that will be achieved when running Forward, and is by default parameterized such that ∼ 1/1000 random sequences is expected to pass the filter. This Viterbi filter is, in turn, preceded by an even simpler filter that compares the query to the target using a scoring scheme with further reduced precision (8-bit integers) and a variant of the model that does not allow for gaps (insertions or deletions). In the context of nucleotide alignment, the HMMER search tool nhmmer [15] calls this stage the Single Segment ungapped Viterbi (SSV) algorithm. SSV produces a rough approximation of the score that will be produced when computing the full Forward alignment. The approximation is not particularly accurate, but is close enough to be generally useful; empirical evidence [35] suggests that nearly all matches that are reported by an unfiltered Forward implementation will also survive the Viterbi filter at *p* ≤ 0.001 and the SSV filter at *p* ≤ 0.02 [35]. HMMER3 parallelizes its SSV implementation using 16-way striped SIMD vector parallelization [36] SSE instructions. Windows of sequence around any threshold-passing region are then subjected to a full Viterbi alignment. Any regions that pass this filter are then aligned with the full Forward algorithm. In total, SSV run time is ∼ 30X faster than the Forward implementation, and is thus key to HMMER’s speed. Even so, SSV accounts for ∼ 70% of HMMER’s run time in common uses, and is thus a performance bottleneck.

### Algorithm 1 The Single Segment Ungapped Viterbi Algorithm

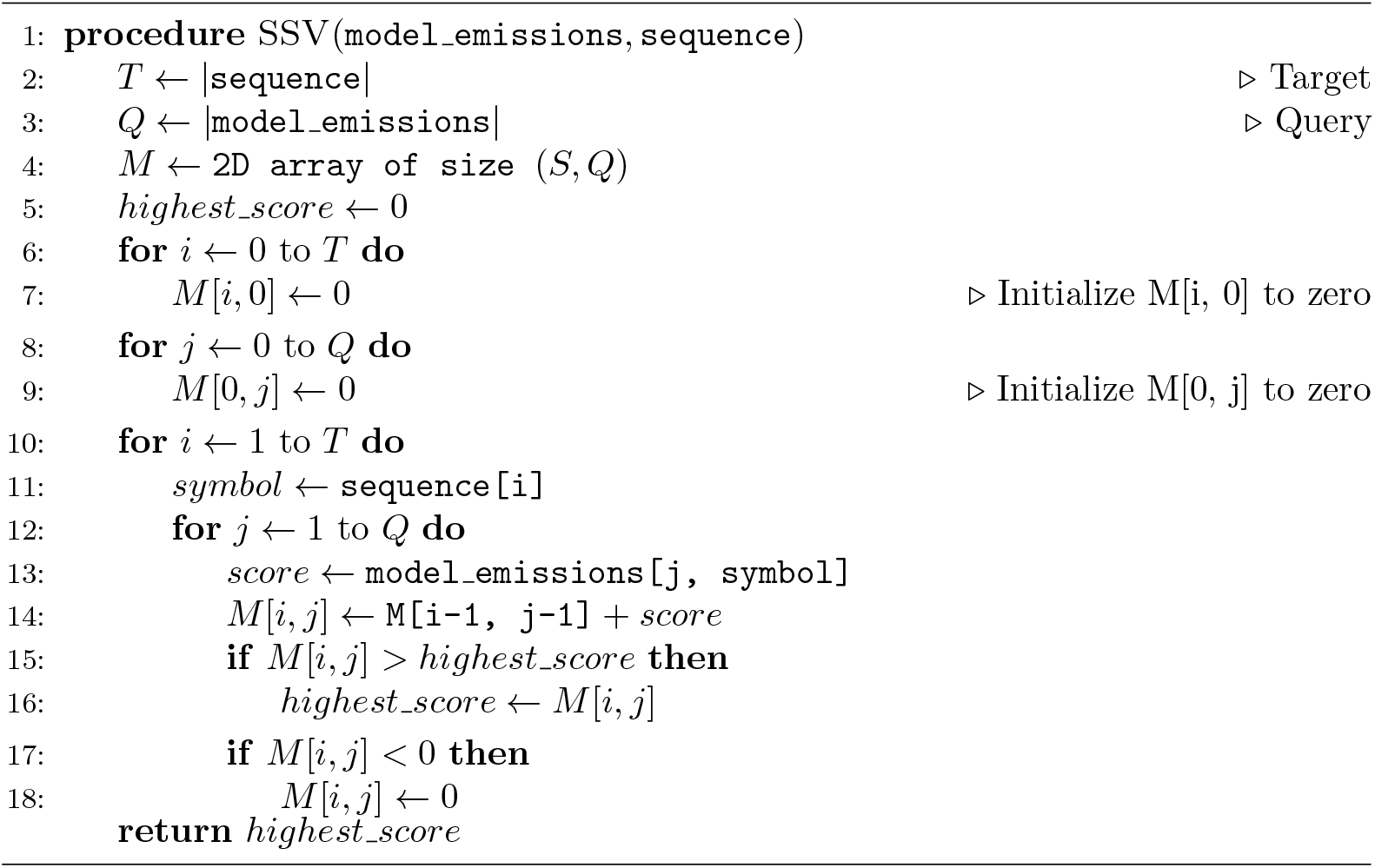

## Hardware Acceleration

An alternative approach to improving the runtime performance of the sequence homology search algorithms has been to employ hardware acceleration. Numerous FPGA hardware accelerators for the Smith-Waterman algorithm [38–41] take advantage of the natural parallel processing ability of configurable hardware to align protein sequences to each other. ClawHMMER [42] was the first work to accelerate the pHMM Viterbi algorithm using Graphics Processing Units (GPUs). CUDAMPF [43] later used GPUs to accelerate ungapped Viterbi, gapped Viterbi, and Forward/Backward in HMMER. Other works have shown that Field Programmable Gate Arrays (FPGAs) can be used to further improve the pararallel performance of dynamic programming-based sequence homology algorithms. FPGAs are hardware devices that can be configured by Hardware Description Language (HDL) code to implement arbitrary digital logic circuits. Unlilke CPUs and GPUs, FPGAs have no prestructured computational architecture and can be optimized for a specific computational task. For example, an FPGA can be used to implement a chain of Processing Elements (PEs) that feed their outputs directly into the inputs of the next PE in the chain. This hardware structure is called a systolic array, and can be used to efficiently pass data along in a highly parallel computational environment.

In serial implementations of sequence alignment, as in Smith-Waterman or in Viterbi and Forward with the standard model (Fig 2A), it is common to perform calculations in row-major order, so that all calculations on one row are performed before moving on to the next row. Straightforward parallel implementations in which a batch of cells on one row are computed concurrently suffer from a data dependency pattern in which each cell depends on the previous cell in the row as well; multiple cells cannot be computed concurrently without either speculative calculation techniques as in [44] or parallelizing computation along the dynamic programming matrix’s anti-diagonal [45].

**Fig 1.**
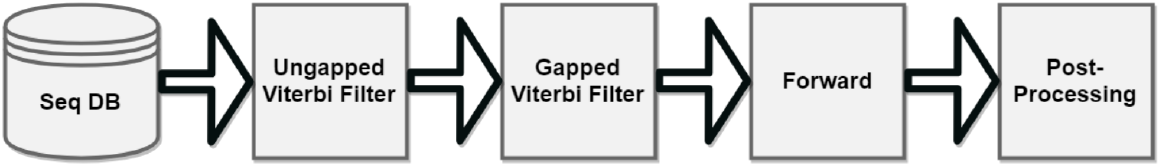
HMMER Pipeline. The major stages of the HMMER pipeline. The Ungapped Viterbi, Gapped Viterbi, and Forward stages of the pipeline function as filters, reducing the number of queries that are passed onto subsequent stages.

**Fig 2.**
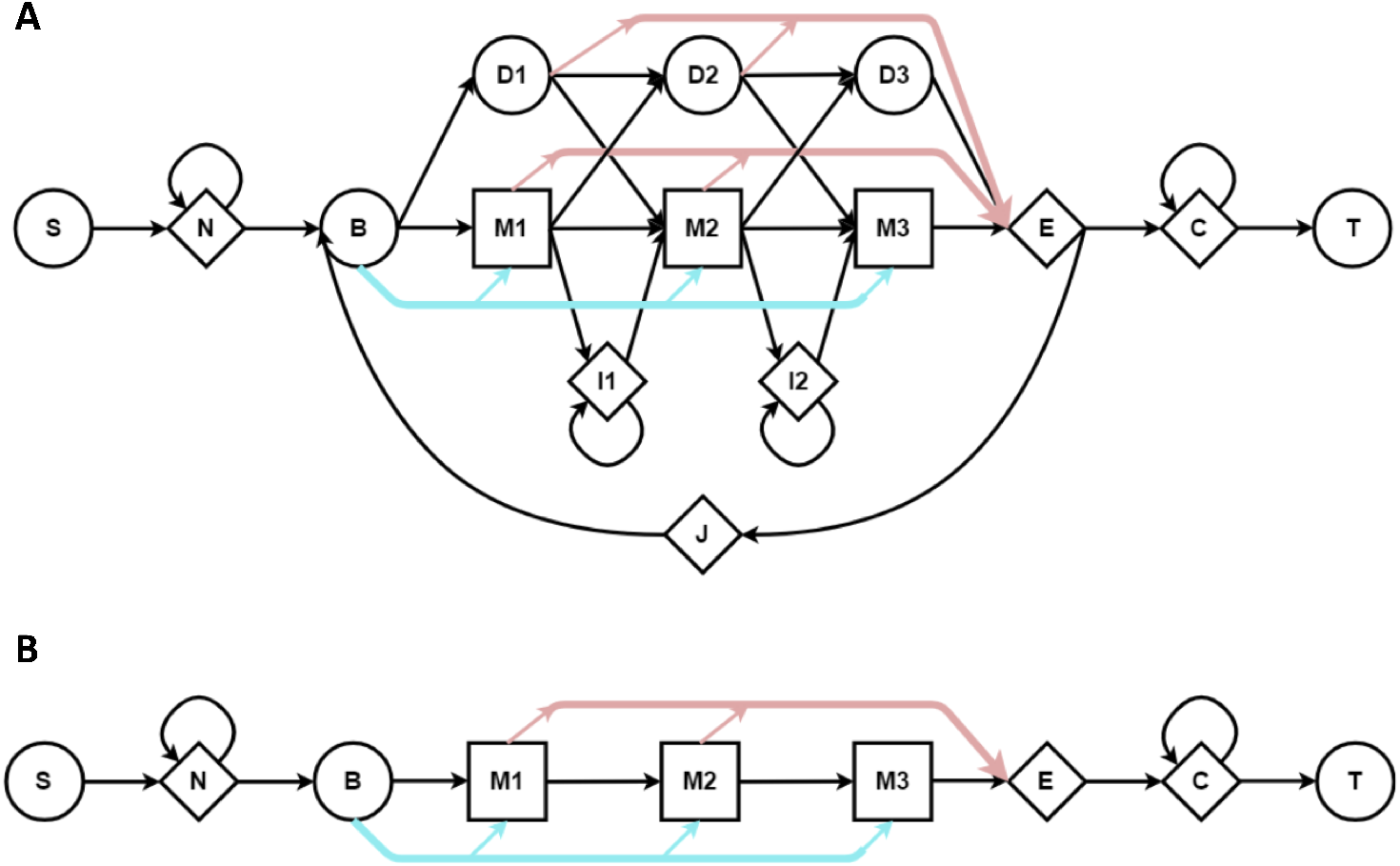
The standard and SSV state models for HMMER3 pHMMs. In (**A**), the core model utilized by HMMER3 Viterbi and Forward algorithm consists of states for observed positions in the modeled family (M), states for insertions relative to those positions (I), and silent states corresponding to the loss or deletion of those positions (D). The Jump state (J) enables a match between the target genome sequence and two disconnected (or even repeated) regions of the aligned model. The other states and path-skipping edges (blue/red) are required for proper scoring statistics [37]. (**B**) shows the reduced model utilized by the SSV algorithm implemented in HMMER3 and HAVAC, in which I, D, and J states are removed; this reduces data dependencies between cells of the Dynamic Programming matrix, and corresponds to alignments between query and target that contain only consecutive aligned positions.

**Fig 3.**
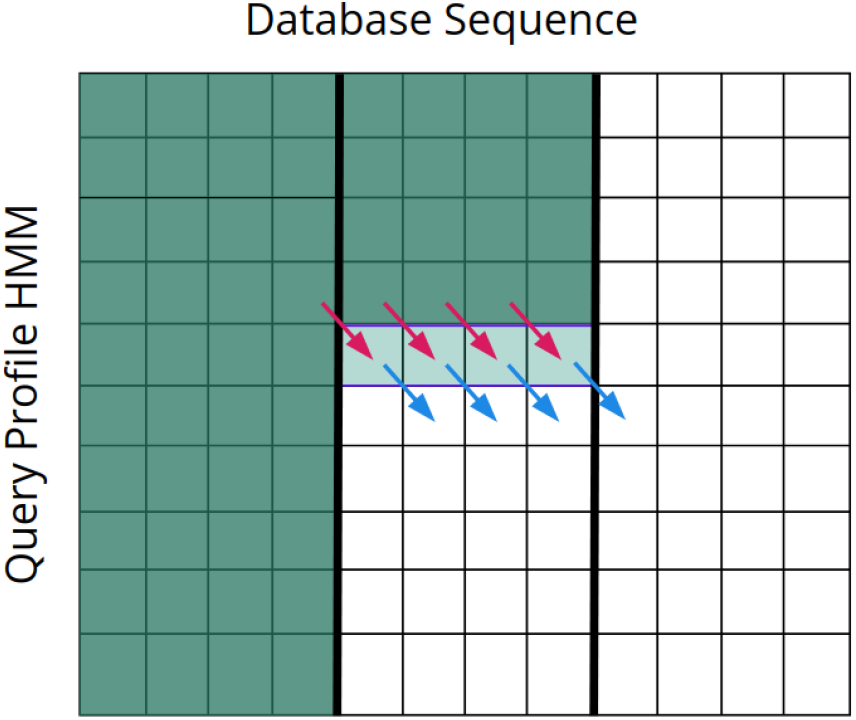
SSV dynamic programming matrix. The Dynamic Programming matrix for the SSV model. Here, the sequence is comprised of 3 segments of length *N* = 4. The light colored row shows the cells being computed this cycle. The fuchsia arrows indicate the data from the cells *M*(*i*–, *j* –;). The blue arrows indicate the cells that will use the data generated this cycle to compute *M*(*i* +1, *j* +1)

**Fig 4.**
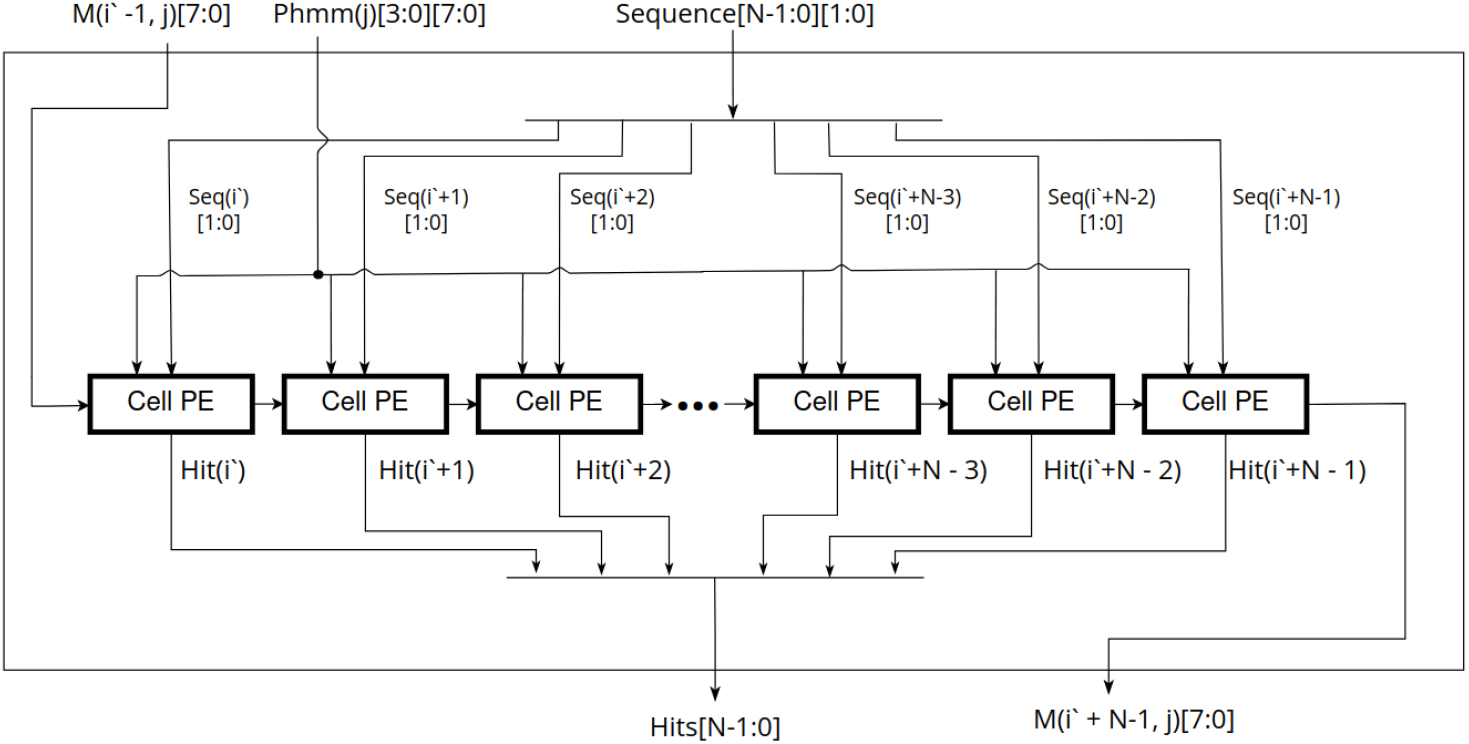
Systolic array of *N* cell PEs. Each cell PE receives the score that was computed in the previous PE on the previous cycle. All cells use the same pHMM vector on a given cycle, and advance to the next vector on the subsequent cycle. Each cell uses a different symbol from inside the contiguous sequence segment to select the correct emission score for the PE’s corresponding DP matrix cell.

**Fig 5.**
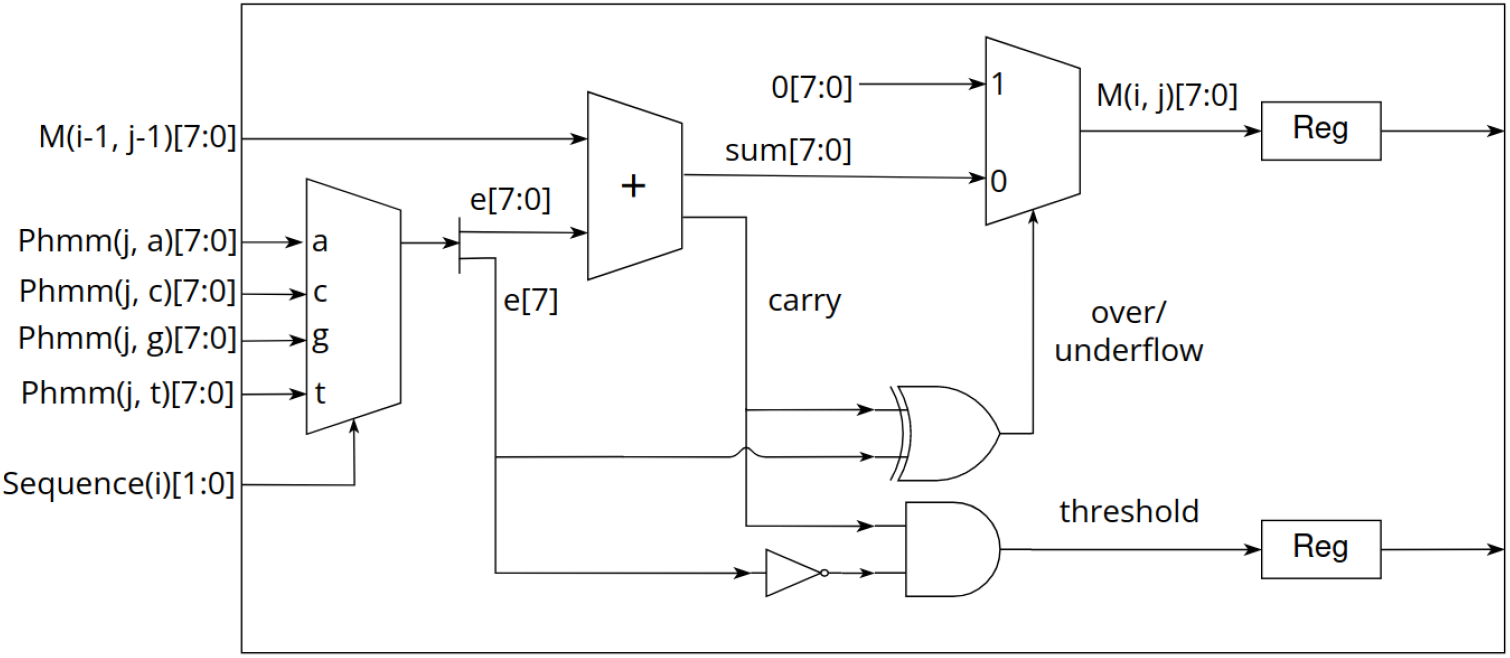
Logic Diagram of the Cell PE. The pHMM vector’s four 8-bit scores corresponding to each nucleotide are de-multiplexed by the sequence symbol, and then added to the score from the previous cell PE. Simple bitwise operations then determine if an overflow or underflow occurred, and reset the score or report a hit accordingly.

**Fig 6.**
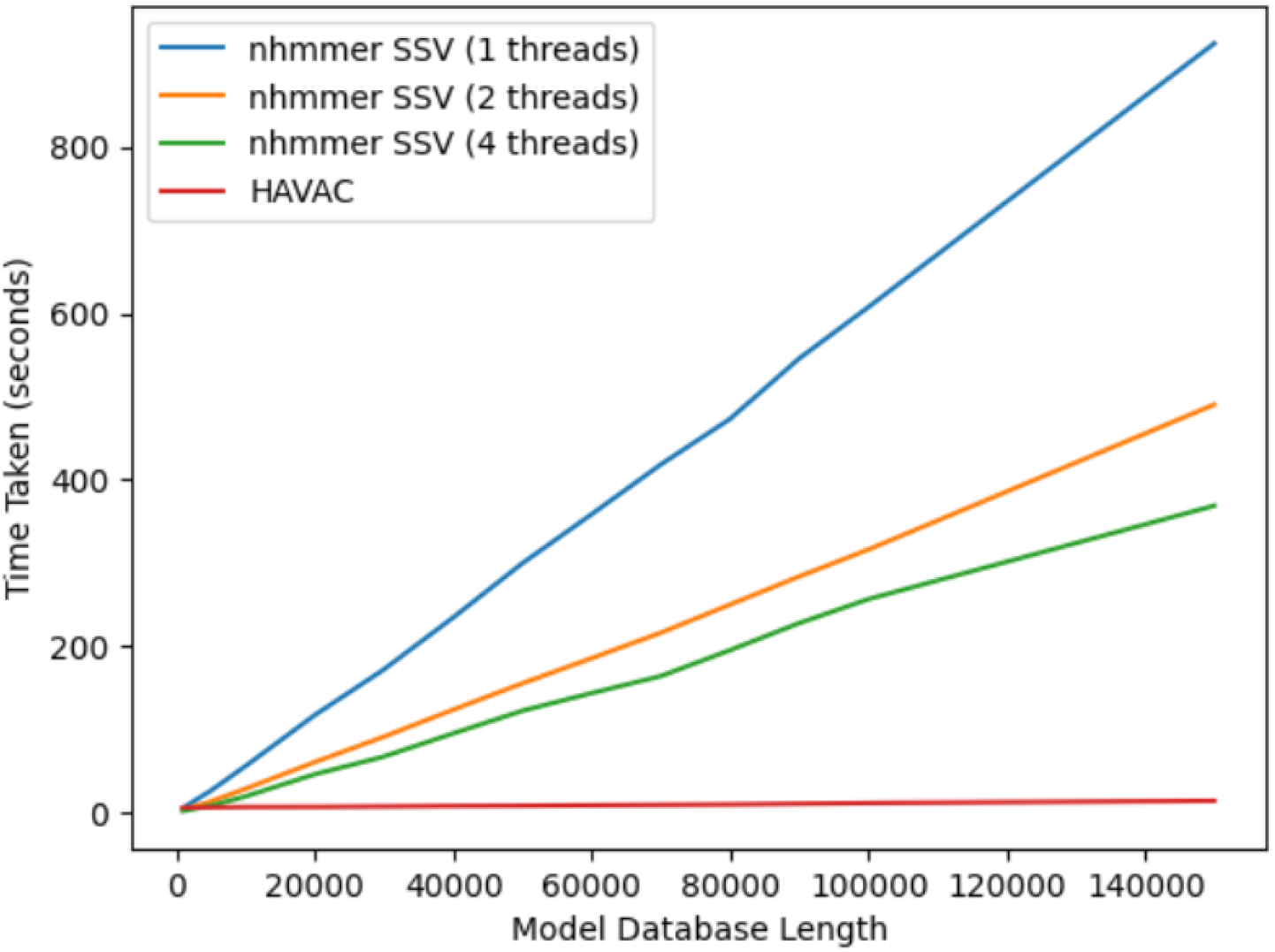
Runtimes of HAVAC and nhmmer SSV at various numbers of threads

An additional dependency challenge is presented by the existence of the J state, which may increase search sensitivity by allowing multiple passes through the core model. The inclusion of the J state results in a data dependency between each cell in the DP matrix and every cell corresponding to the previous sequence character, and as a result the maximum score from a given row must be identified before any cells in the subsequent row can be computed. Maddimsetty et al. [46] implement a two-pass hit detection algorithm by appending a duplicate copy of the model to the end of the model in lieu of the J state. This effectively allows two separate homologous areas to accumulate score as if there is a single-use J state loop. Oliver et al. [47] implements Viterbi on the full HMMER model including the J state, opting to parallelize over multiple discrete sequence/model pairs instead of along a single DP matrix anti-diagonal.

Abbas et al. [48] goes beyond Viterbi to implement a single accelerator that runs both a Multiple-Sequence Ungapped Viterbi (MSV) filter and Viterbi on the same hardware. The MSV filter is similar to SSV, but it retains the HMMER pHMM J state. Abbas et al. implement the MSV filter with a max-reduction tree across a static number of cells from previous model states, allowing the accelerator to approximate the J state in a reasonable amount of parallelizable work, at the cost of a potential drop in sensitivity.

Another strategy is to directly follow the philosophy of the HMMER filtering pipeline, and implement an FPGA accelerator focused entirely on the SSV model – this model removes the J state as well as all insert and delete states, leaving a minimal dependency map. Here, we introduce such an accelerator, the Hardware Accelerated single-segment Viterbi Additional Coprocessor (HAVAC), a hardware accelerator that implements SSV as a standalone nucleotide sequence homology filter. HAVAC is designed to exist as a standalone nucleotide sequence homology filter that can be incorporated into a pHMM alignment pipeline in which SSV filter is run on the FPGA accelerator while downstream Viterbi and Forward alignment algorithms can be run simultaneously on the host CPU. To our knowledge, this is the first FPGA hardware accelerator that implements SSV for sequence homology search filtering. The relative computational simplicity of the SSV model allows for better parallel performance than its more intricate counterparts. We also present a method for scaling a profile HMM’s emission scores to leverage an 8-bit full adder’s carry bit to check if a model’s state score passes a significance threshold without requiring an 8-bit comparator. HAVAC returns the (*i, j*) pairs of DP matrix cells that pass the given threshold to allow later stages in a sequence homology search pipeline to localize search around areas of likely homology. HAVAC supports fasta-formatted sequence files and HMMER3-formatted model files. HAVAC is open-source licensed under BSD-3 and is available at https://github.com/TravisWheelerLab/HAVAC.

## Methods

The HAVAC hardware accelerator was implemented on a Xilinx Alveo U50 FPGA accelerator card. The Alveo U50 contains a custom-built Ultrascale+ FPGA with approximately 874K lookup tables (LUTs) and supports two 4GiB banks of High Bandwidth Memory (HBM). The HAVAC hardware design was implemented using Vitis High Level Synthesis (HLS), a tool for synthesizing designs from compliant C/C++ codebases along with FPGA-specific #pragma instructions. Vitis HLS can allow for significantly faster development compared to traditional hardware description languages (HDLs), at a small cost to implemented resource utilization and performance [49]. The HAVAC host (CPU) driver code was implemented in C++ using the Xilinx Runtime Library (XRT).

### Profile HMM scores

As in the nhmmer SSV filter, HAVAC computes the scores of the Dynamic Programming (DP) matrix using 8-bit integer emission sores. For a given pHMM, a threshold score *t* is generated representing the minimum score required for a target database sequence to pass the requested P-value target (default: *P≤*0.02) using the pHMM’s pre-computed gumbel distribution parameters [37] with adjustments to account for SSV’s removed state transitions. HAVAC uses the threshold to generate a scaling factor *τ* on the pHMM’s emission scores. The purpose of *τ* is to reproject the emission scores such that an accumulated score passes the query-specific threshold if and only if the score reaches 256.

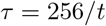

The purpose of using *τ* to re-project the emission scores is twofold. The primary reason is to simplify the hardware accelerator’s cell Processing Elements (PEs). By enforcing a threshold score of 256, the cell PEs can check their score against the threshold by using an 8-bit full adder’s carry bit, eschewing the need for a full 8-bit comparator to check for hits. As a result, cell PEs require fewer resources to implement, which allows more PEs to fit into the hardware to improve parallelism. As an added bonus, using *τ* to reproject the emission scores allows for better use of the 8-bit integer space, allowing for slightly more precision in the emission scores when compressed down to 8-bit integers.

The HMMER3 pHMM format stores emission scores as negative log-likelihood values, and these scores must be converted to bits before being projected using *τ* . To convert a negative log-likelihood score s to a *τ* -reprojected score *s*^*′*^ in bits, the score is represented as a single-precision floating point value and extracted from negative log-likelihood space, divided by the background distribution of equal probability for each of 4 nucleotides, converted to bits, and then multiplied by *τ* .

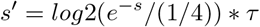

The above equation involves performing exp() and log() function calls on each emission score, which are slow on modern hardware. We eliminate the need to perform these expensive functions by simplifying as follows.

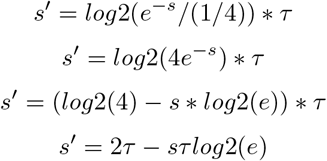

In the resultant equation, the terms 2*τ* and *τlog*2(*e*) can be computed as constants over all match states of the model; this means that all match states can be computed with a single multiplication and subtraction. Each reprojected emission score s is then rounded to the nearest integer, and cast to signed 8-bit integers. Each individual pHMM in the input .hmm file is projected in this way using the same P-value and the data are appended together and written to the FPGA’s HBM memory banks.

### SSV Processor Design

Define *M* (*i, j*) as the score of the maximum scoring ungapped Viterbi alignment ending at the *i*th letter of the sequence and the *j*th position of the pHMM. For a sequence of length *L* and a pHMM of length *K*, this requires computation of *LK* cells. The HAVAC kernel on the FPGA is comprised mainly of a systolic array of *n* cell PEs that each individually compute a single cell of the DP matrix each cycle. If *L > n*, then the sequence is procedurally broken into length-*n* segments. Each cell PE in the systolic array uses a different letter from the sequence to find the match score for the current pHMM position and adds it to the score computed by the previous cell PE on the previous cycle. Initially, the systolic array computes the cells *M* (1, 1) to *M* (*n*, 1) in the DP matrix. The model position is then incremented and the systolic array computes *M* (1, 2) to *M* (*n*, 2), and so on until a full pass through the pHMM has been completed. If *L > n*, then a new segment of the sequence is loaded into the cell PEs and another pass through the pHMM begins. These passes through the pHMM compute columns of width *n* through the DP matrix until the entire matrix has been calculated. Sequences that are not a multiple of *N* in length are padded with random data to reach that target; SSV matches to these random sequences will be rare, and are easily filtered out by host-side driver.

For any given sequence segment, the leftmost cell of the systolic array of cell PEs requires input scores from the previous sequence segment column (except the first segment, which requires inputs of 0 for each cell in the first column of the matrix). Likewise, the scores generated by the rightmost cells in the segment column must be used as inputs for the next segment column of the matrix. A hardware FIFO structure is used to enqueue the outputs of the rightmost cell in a segment column, and dequeue scores as needed by the next segment column. Since hardware FIFOs have a strict maximum capacity, this acts as a limiting factor on the total length of phmm vectors that HAVAC can support. HAVAC supports a maximum of 1,048,576 (1024*1024) model positions for a single SSV query. This is large enough to hold the entirety of the Rfam database of RNA families [50](4108 models) twice over.

### Hardware Cell Processing Elements

The HAVAC cell PE is designed to minimize the resources required to calculate the score at a given cell. The cell’s sequence symbol *c* is used as the *select* on the current pHMM vector *P* (*j*) to obtain the cell’s signed 8-bit pHMM emission score. This value is then summed with the score computed by the previous cell PE on the previous cycle, *M* (*i* 1, *j* 1), to obtain an unsigned 8-bit intermediate sum value *T* (*i, j*) and a carry bit from the 8-bit full adder. This carry bit is then used along with the sign bit from the emission score to determine if the threshold has been passed and if the final cell’s result, *M* (*i*–, *j*–), should be reset back to zero. There are two situations where the result of the summation should be discarded: if a negative match score was added to *M* (*i*–, *j* –) (the carry bit was clear), or if a positive emission score was added to *M* (*i*–, *j*–) (the carry bit was set). In the first case the result underflowed the unsigned 8-bit result, and in the latter case the result overflowed the maximum representable score of 256. In this second case, the given cell has passed the score threshold and should be reported.

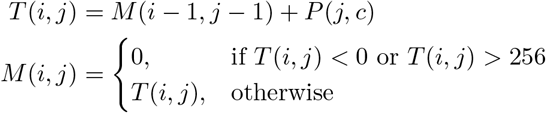

### Hit Reporting

HAVAC reports any cell that passes the 256 threshold as an (*i, j*) position pair in the sequence and pHMM. HAVAC finds these (*i, j*) pairs by taking the threshold bit from all *N* cell PEs and systematically partitioning the threshold hit bits to find and report any that were set for a given cycle. First, all *N* threshold hit bits are *bitwise-or* reduced; if the result is 1, the threshold hit bits contain at least 1 cell that passed the threshold. In this case, the threshold hit bits are enqueued to a hardware FIFO along with the current pHMM position and the current sequence segment index. Then, in concurrently running processes, the threshold hit bits are partitioned into 16 contiguous bit ranges. HAVAC iteratively *bitwise-or* reduces each of these 16 bit ranges to determine if a hit was in any of the ranges. If a bit range contained a hit, the bit range is enqueued to a subsequent hardware FIFO along with the rest of the position metadata and the index of the bit range. This process continues, further dividing the threshold hit bits until it represents a single bit. At this point, the partition indices along with the sequence segment index can uniquely identify the exact sequence position of the hit. The hits are then written to the FPGA HBM memory where they can be read by the host to extract the (*i, j*) pairs after the HAVAC hardware accelerator finishes.

Because the hit report partitioning runs concurrently to the main systolic array, and the hardware doesn’t know a priori how many hit reports will be written for an invocation of the hardware accelerator, an extra terminator bit t is included along with the threshold hit ranges and positional data and is normally cleared. On the final cycle of the matrix computation (the final pHMM position of the final sequence segment index), the full threshold hit bit range is enqueued to the partitioning system even if it doesn’t contain any hits, along with a set terminator bit. Each subsequent partitioning tier then passes along a threshold bit range with a set terminator on the final (16th) section of the bit range, and then deactivates. Once all hit reports are written to memory and the final partitioning tier deactivates, the number of hits is written to memory and the hardware accelerator completes its operation.

### Accelerator Synthesis and Driver

Synthesis of the HAVAC hardware accelerator design was performed inside the Xilinx Vitis HLS tool. Implementation was performed by the v++ compiler tool. Of the 8GiB of HBM available, 4GiB is used to store sequence data. All sequence characters are represented as 2-bit encodings (ambiguity characters are replaced by random letters, which may result in rare cases of falsely passing the filter; these are quickly filtered out by the downstream driver). HAVAC supports target sequences up to an aggregated length of 16Gbp. 3.5GiB is allocated for reporting the 8 byte (*i, j*) pairs for hits. Therefore, a maximum of 469,762,048 hits can be reported for any query. HAVAC was implemented with 12288 cell PEs at a clock speed of 144.5 MHz.

The HAVAC driver software library allows for simple control over the hardware accelerator. Input data is provided to the driver as a fasta file and a HMMER3 model file and is then preprocessed and loaded onto the board’s HBM memory banks. Users can then begin the SSV computation either synchronously or asynchronously. Once the hardware accelerator has finished, hit data can be read from the device as global sequence and model positions. Using these global positions, the HAVAC driver references the fasta and model files to locate which sequence and model the hits resides in, and the local positions in the sequence or model the hit refers to. These sequence and model positions can then be used by other algorithms in a sequence homology pipeline as positions of likely homology. Because of the nature of HAVAC being a dedicated co-processor, SSV calculation may be performed concurrently to these other algorithms, further reducing the processing time for large queries that can be broken down into smaller batches.

## Results

We performed benchmarking experiments on a compute system with an Intel Xeon X5570 CPU @ 2.96GHz and 114 GB of memory. The test data was stored on an Intel 240GB SSD. Tests were performed by searching human chromosome 22 against query databases generated from subsets of the Rfam family model database. Query datasets were generated by selecting models from Rfam up to a specified number of model positions. The HAVAC hardware accelerator was implemented on a Xilinx Alveo U50 Data Center accelerator card. FPGA routing and implementation typically uses only a fraction of the full board capacity – HAVAC’s final hardware implementation utilized 48.94% of the device’s hardware lookup tables (LUTs), 21.53% hardware register, and 35.94% Block RAMs. HAVAC’s runtime was timed as 4 discrete sections; (i) software data allocation and configuring the FPGA with the Xilinx hardware device binary (.xclbin) file, (ii) file I/O for the sequence and pHMM files, data preprocessing, and loading these data onto the device’s HBM banks, (iii) runtime of the HAVAC SSV kernel, and (iv) reading the generated hits from the device memory and resolving the global (*i, j*) pairs to local sequence and model positions. It is common to describe the performance of dynamic programming algorithms in terms of billions of cells updated per second (GCUPS). HAVAC’s performance was measured to be 1739 GCUPS using the runtime of the kernel from step (iii). This performance represents a realization of 98% of the theoretical maximum speed achievable with 12,288 cell PEs processing a cell every cycle at 144.5 MHz. In comparison, HMMER3 contains a highly optimized CPU implementation of MSV that utilizes 16-way striped SIMD parallelization reported to reach 12 GCUPS on a single thread running on a 2.66 GHz Intel Gainestown X5550 CPU [27]. The nhmmer SSV matrix calculation on our tests system averaged 7.6 GCUPS when single-threaded, and 18.9 GCUPS with four threads. We tested nhmmer with 8 and 16 threads, but saw performance that was nearly identical to 4 threads; this is consistent with reports that HMMER3 becomes I/O bound at low thread count and does not benefit from additional threads past this point [51, 52]. The HAVAC kernel represents a 227x matrix calculation speedup over nhmmer with one thread and a 92x speedup over nhmmer with 4 threads. Table 2 shows the performance in terms of GCUPS for HAVAC and previous publications of Viterbi-family algorithms implemented on FPGAs and GPUs.

**Table 1.**
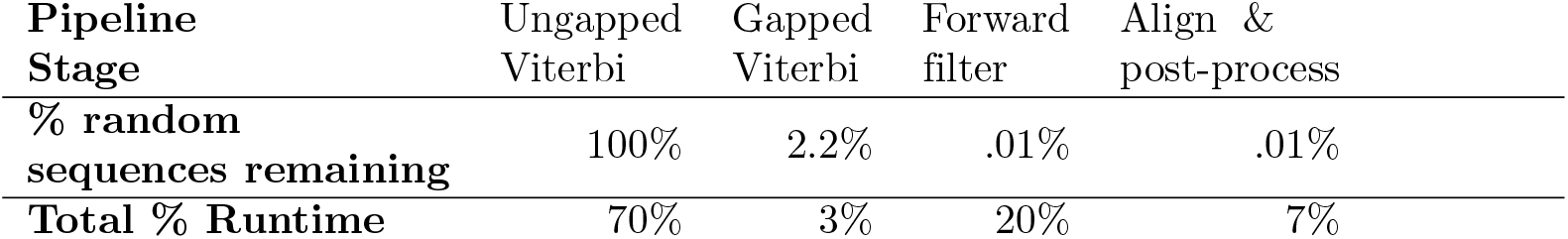
The major stages of the HMMER3 pipeline, the percentage of random sequences that will pass a given pipeline stage, and the approximate runtime takes by each stage as a percentage of total runtime.

| Pipeline Stage | Ungapped Viterbi | Gapped Viterbi | Forward filter | Align & post-process |
| --- | --- | --- | --- | --- |
| % random sequences remaining | 100% | 2.2% | .01% | .01% |
| Total % Runtime | 70% | 3% | 20% | 7% |

**Table 2.** Fastest reported FPGA and GPU accelerator speeds for Viterbi-family algorithms on biological data from the literature.

| Implementation | Algorithm | Year | Maximum Reported GCUPS |
| --- | --- | --- | --- |
| Jacob et al. [53] | Viterbi | 2007 | 10.6 (best case estimate) |
| Walters et al. [54] | Viterbi | 2007 | 0.7 |
| Benkrid et al. [55] | Viterbi | 2008 | 9 |
| Oliver et al. [56] | Viterbi | 2008 | 2.1 |
| Md Isa et al. [57] | Viterbi | 2012 | 11.8 |
| Abbas et al.[58] | MSV | 2015 | 81 |
|  | Viterbi | 2015 | 3.6 |
|  | MSV + Viterbi | 2015 | 45.7 |
| Jiang & Ganesan [59] | SSV (GPU) | 2016 | 440 |
|  | MSV (GPU) | 2016 | 277 |
|  | Viterbi (GPU) | 2016 | 14.3 |
| **HAVAC** | SSV | 2023 | 1739 |

HAVAC’s performance was also measured as a complete SSV filter including all steps from initial FPGA configuration to resolution of hit positions. These times were compared to nhmmer’s SSV filter using 1, 2, and 4 threads with the subsequent stages of the HMMER pipeline disabled. At small query sizes, the hardware configuration and PCI-e data transfer times dominate run time, resulting in poorer performance than nhmmer. HAVAC and nhmmer with a single thread had similar runtime with a query length of 1000 model positions (representing 6 models), and nhmmer with 4 threads was similar with a query length of 3000 model nodes (representing 16 models). As the size of the DP matrix grows, the hardware kernel’s performance becomes the dominating factor – when querying the entire Rfam database (4108 models) against human chromosome 22, HAVAC is 65x faster than nhmmer with 1 thread.

Power utilization of the HAVAC hardware accelerator was approximated using the Xilinx Board Utility (xbutil) tool. Voltage and Amperage values were reported for the PCI-express interface and on-board power rails. These mV and mA values were multiplied to find the watts used, and them summed to total energy use estimate of 29 watts, 31% of the 95 watts of the Thermal Design Power (TDP) of the Intel Xeon X5570 in our test system [60]. This TDP represents the average energy usage when all cores are under full load.

## Conclusion

In this work, we have described a new implementation of HMMER’s SSV filter algorithm on configurable hardware with excellent runtime characteristics. The limited data cell dependencies of SSV allow design of cell PEs that consume very limited resources. Considering that SSV represents 72% of nhmmer’s runtime, a 31x speedup over the current SSV implementation would result in a ~3.5x improvement on the pipeline overall when given 4 threads.

At 50% LUT and 22% register utilization, HAVAC could likely be reimplemented to increase the number of cell PEs in the design. Attempts were made to increase the number of cell PEs from 12288 to 16384, but the Xilinx v++ implementation stage was unable to generate the design. This is not unreasonable – as utilization grows, FPGA implementation becomes significantly more difficult as signals must be routed around increasingly more congested sections of the design, and may not be able to reach their destination in a reasonable amount of time. Smaller increases to the number of cell PEs are likely possible, although such changes would require modifications to the hit report partitioning system. Importantly, the Alveo U50 represents the lower end of the Xilinx data center accelerator cards, and a mid-range of FPGA accelerators in general, and it is reasonable to expect even faster HAVAC implementations on more powerful hardware.

It is important to remember that the SSV algorithm is a fast approximation of the Viterbi algorithm, but has limited sensitivity by itself. HAVAC efficiently filters to find areas of potential homology between sequences and probabilistic models, but is not an effective tool without a downstream pipeline of more accurate algorithms. As such, HAVAC is only an effective tool in conjunction with the rest of a bioinformatics pipeline. Future work would necessarily involve combining HAVAC with CPU-based downstream Viterbi or Forward algorithms to make a tool that would be applicable to the overall task of genome annotation.

## Acknowledgements

We thank George Vacek who piqued our interest in FPGA acceleration of pHMM search while at Convey Computing. We are grateful to the Computer Architecture Group (and especially Farzad Fatollahl-Fard) at Lawerence Berkeley National Labs (LBNL) for providing access to a development FPGA system early in the project. We thank Xilinx for providing a free Alveo U50 FPGA accelerator platform through their Xilinx University Program. Additionally, Patrick McInerney for suggestions for simplifying the equation for profile HMM reprojection. We also gratefully acknowledge the computational resources and expert administration provided by the University of Montana’s Griz Shared Computing Cluster (GSCC).

